# A mitochondria-targeted antioxidant and a thyroid hormone affect carotenoid ketolase gene expression and bill redness in zebra finches

**DOI:** 10.1101/2020.01.14.905745

**Authors:** Alejandro Cantarero, Pedro Andrade, Miguel Carneiro, Adrián Moreno-Borrallo, Carlos Alonso-Alvarez

## Abstract

Conspicuous ornaments in animals can evolve to reveal individual quality when their production/maintenance costs make them reliable as signals or if their expression level is intrinsically linked to quality by some unfalsifiable mechanism (quality indices). The latter has been mostly associated with traits constrained by body size. However, red ketocarotenoid-based coloured ornaments may also have evolved as quality indices because their production could be closely linked to individual metabolism and, particularly, to the cell respiration at the inner mitochondrial membrane (IMM). This mechanism would supposedly not depend on resource (yellow carotenoids) availability, thus discarding allocation trade-offs. A gene coding for a ketolase enzyme (CYP2J19) responsible for converting dietary yellow carotenoids to red ketocarotenoids has recently been described in birds. It is not known, however, if this ketolase is involved in mitochondrial metabolism and if its expression level and activity is resource independent. Here, we manipulated the metabolism of captive male zebra finches by an antioxidant designed to penetrate the IMM (mitoTEMPO) and a thyroid hormone (triiodothyronine; T3) with known hypermetabolic effects. The expression levels of a ketocarotenoid-based ornament (bill redness) and *CYP2J19* were measured. MitoTEMPO downregulated *CYP2J19* expression, supporting the mitochondrial involvement in ketolase function. T3 also reduced *CYP2J19* expression, but at an intermediate dosage, this effect being buffered by mitoTEMPO. Bill redness seemed to show a similar interacting effect. Nevertheless, this faded when *CYP2J19* expression level was controlled for as a covariate. We argue that the well-known mitoTEMPO effect in reducing mitochondrial reactive oxygen species (ROS) production (particularly superoxide) could have interfered on redox signalling mechanisms controlling ketolase transcription. High T3 levels, contrarily, can lead to high ROS production but also trigger compensatory mechanisms, which may explain the U-shaped effect with dosage on *CYP2J19* expression levels. Bill *CYP2J19* expression values were also positively correlated to redness and circulating substrate carotenoid levels. Nonetheless, treatment effects did not change when controlling for blood carotenoid concentration, suggesting that resource-availability dependence was irrelevant. Finally, our findings reveal a role for thyroid hormones in the expression of carotenoid-based ornaments that has virtually been ignored until now.

## Introduction

The animal signalling theory proposes that traits involved in animal communication, such as many conspicuous ornaments and songs, evolve (1) due to production/maintenance costs that prevent cheating by low-quality individuals (Grafen 1990) or (2) because the level of expression of the trait directly reveals individual quality (i.e. they cannot be faked; Maynard Smith & Harper 2003). The first type of trait has often been defined as “signals” (or “handicap signals”), whereas the second type of trait has been defined as “indices” (e.g. Johnstone 1995; Weaver, Koch & Hill 2017). The expression level of sexual “signals” should in some way correlate positively with reproductive success, but negatively to survival, due to some direct or indirect costs of trait production/maintenance. Instead, an “index” should positively correlate to both reproductive success and survival (i.e. no trade-off should exist; Vanhooydonck *et al*. 2007) and is supposedly cost-free (but see Biernaskie, Grafen & Perry 2014). Most examples of indices have been associated with traits (e.g. deer antlers or calling rates) that depend or are positively correlated to body size (Maynard Smith & Harper 2003; Reby & McComb 2003).

However, a new quite different type of index has been proposed in the form of conspicuous colourations produced by red carotenoid pigments (Hill 2011; Weaver *et al*. 2017). Red carotenoid-based ornaments are present in many vertebrates. They have attracted much attention from evolutionary ecologists as its proximate production mechanisms are intriguingly complex (e.g. McGraw 2006). Such complexity is, however, becoming to be disentangled (e.g. Johnson & Hill 2013; Lopes *et al*. 2016; Mundy *et al*. 2016), providing material to propose new evolutionary hypotheses in the signalling theory framework (Hill 2011). The underlying question is why these mechanisms have promoted the selection of red ornaments as reliable information transmitters.

Carotenoids are molecules whose chromophore generates yellow to red colourations (Britton 2008). The animal organism cannot synthesize them from any non-carotenoid substrate, but some carotenoids can be transformed into others by enzymatic reactions (Stradi *et al*. 1997; McGraw 2006; García-de Blas, Mateo & Alonso-Alvarez 2016). This is the case of red ketocarotenoids (e.g. astaxanthin, canthaxanthin) obtained from dietary yellow-to-orange carotenoids (e.g. lutein, zeaxanthin). This mechanism has been well studied in birds (McGraw 2006). Many aquatic bird species can easily obtain red ketocarotenoids from their food and directly allocate them to ornaments without transformation because ketocarotenoids are abundant in aquatic invertebrate prey (e.g. crustaceans; McGraw 2006). However, among terrestrial birds, ketocarotenoids are often scarce in food (particularly in vegetal food) and molecular mechanisms allowing to transform yellow to red pigments have evolved (McGraw 2006). Currently, the only candidate gene implicated in this transformation and well-supported by molecular studies is *CYP2J19* (Lopes *et al*. 2016; Mundy *et al*. 2016; Twyman *et al*. 2016; 2018). This is a member of the cytochrome p450 family of enzymes, most of its members being involved in the metabolism of toxicants but also in hormone synthesis (Tompkins & Wallace 2007).

Before the discovery of *CYP2J19*, some researchers argued that the candidate enzyme should be linked to mitochondrial activity (Hill 2011; Johnson & Hill 2013). They argued that the molecular similarity between some ketocarotenoids and ubiquinone, which is a key antioxidant involved in the cell respiratory chain, implies that the enzyme could be part of the ubiquinone enzymatic biosynthesis pathway (Johnson & Hill 2013). The transformation would, hence, be made in the inner mitochondrial membrane (IMM) probably sharing a biochemical pathway with cell respiration. The theoretical link to the mitochondria metabolism led these authors (see Hill 2011; Hill & Johnson 2012) to propose that red ketocarotenoid based ornaments evolved as “indices” of individual quality as they would be tightly linked to basic metabolic pathways such as cell respiration.

The subsequent description of *CYP2J19* could support this hypothesis if the enzyme is indeed placed at the IMM affecting cell respiration. The recent finding of relatively high levels of ketocarotenoids at the IMM compared to other cell fractions is consistent with this scenario (Hill *et al*. 2019). Moreover, the treatment of male zebra finches (*Taeniopygia guttata*) with a synthetic ubiquinone (mitoQ; Smith *et al*. 2003) designed to penetrate into the IMM has been shown to increase bill redness (Cantarero & Alonso-Alvarez 2017), which is a ketocarotenoid-based sexually selected ornament (McGraw & Toomey 2010).

In the present study, we exposed male zebra finches to another synthetic mito-targeted antioxidant, testing, for the first time, potential changes in bill *CYP2J19* expression levels. This experiment allows us to infer if mitochondria metabolism is indeed linked to the candidate gene and if that connection affects trait expression (redness). The antioxidant (i.e. mitoTEMPO: (2-(2,2,6,6-Tetramethylpiperidin-1-oxyl-4-ylamino)-2-oxoethyl)triphenylphosphonium chloride; Dikalova *et al*. 2010) is similar to mitoQ. In both molecules, the antioxidant is joined to a triphenylphosphonium cation (TPP+) specifically designed to penetrate into the IMM (Murphy & Smith 2007). In mitoQ, TPP^+^ is connected to the antioxidant by a 10-carbon alkyl chain (i.e. decyl-TPP^+^). The length of this molecule, however, increases membrane permeability, inhibiting the electron transport chain and rising superoxide radical generation (Reily *et al*. 2013; Trnka, Elkalaf & Anděl 2015; Gottwald *et al*. 2018). Consistent with these negative effects, zebra finches only treated with decyl-TPP^+^ developed paler bills than controls (Cantarero & Alonso-Alvarez 2017). In contrast, mitoTEMPO does not include that linker group and the antioxidant role is played by piperidine nitroxide, which recycles ubiquinol (the reduced ubiquinone form) to ubiquinone (Trnka *et al*. 2008). This lowers mitochondrial superoxide radical concentration (Dikalova *et al*. 2010). Moreover, we have recently found that mitoTEMPO is able to increase ketocarotenoid-based feather redness in males from another bird species (the red crossbill; *Loxia curvirostra*; Cantarero *et al*. 2019 preprint).

Here, we aimed to go further by artificially increasing the level of the most active thyroid hormone (triiodothyronine; T3), testing its impact on both *CYP2J19* expression and redness. Virtually nothing is known about the potential involvement of thyroid hormones in animal carotenoid-based ornaments. However, thyroid hormones control oxygen consumption and have hypermetabolic effects mediated by changes in mitochondria metabolism (Hwang-Bo, Muramatsu & Okumura 1990; Chastel, Lacroix & Kersten 2003; Seifert *et al*. 2008). High blood levels of thyroid hormones are commonly associated with higher oxidative stress, and higher reactive oxygen species (ROS) production, particularly mitochondrial superoxide production (reviewed e.g. in Venditti & Meo 2006; Collin *et al*. 2009; Elnakish *et al*. 2015; Chainy & Sahoo 2019). High oxidative stress may, in turn, exert an inhibitory effect on the activity of CYP enzymatic activity (Zangar, Davydov & Verma 2004). Moreover, high thyroid levels have also been linked to reduced expression of some CYP genes (Honkakoski & Negishi 2000; Kot & Daniel 2011). Accordingly, and also taking into account the precedent results (i.e. Cantarero & Alonso-Alvarez 2017; Cantarero *et al*. 2019 preprint), we hypothesized that mitoTEMPO should increase CYP2J19 activity and bill redness, whereas high T3 levels should decrease them, both treatments interacting perhaps to cancel out their respective effects.

## Material and Methods

### Experimental protocol

Eighty-six male zebra finches were housed in cages placed within an indoor aviary (more details in Supplementary Material, SM). Two birds were housed per cage (0.6 m × 0.4 m × 0.4 m). The pair was divided by a grille hindering physical contact. After an 8-day acclimation period, all the birds were randomly assigned to the treatments. All of them received a subcutaneous silicone implant (OD 1.96 mm, ID 1.477 mm; Silastic®) empty or filled with T3 (3,3′,5-Triiodo-L-thyronine; SIGMA ref. T2822; see also SM). The T3-filled implants were made at different lengths (6 mm, 8 mm and 10 mm) to produce different dosages. Control birds (*n* = 22) received a 10 mm empty implant. The largest length was chosen considering recent studies in Gambel’s white-crowned sparrows (*Zonotrichia leucophrys gambelii* (Perez *et al*. 2016; 2018). Lower dosages were included considering that sparrows are heavier than finches (26 vs. 15 gr approx., respectively). The implants were put in two consecutive days due to time constraints. Thirty-five birds rejected the implant 2-7 days before the end of the experiment (mean ± SD: 3 ± 1.2 days). The treatment distribution among birds losing or maintaining the implant never differs (i.e. by testing both treatments separately or a single eight-level factor all χ^2^ tests: *p*>0.26). The number of days with implant also did not differ among treatments (non-parametric Kruskal-Wallis or Mann-Whitney U’s tests: all *p*-values >0.30). This variable never produced a significant effect (all *p*-values > 0.10) when tested as a covariate in every statistical model, and was therefore removed (see below).

Each implant group (Control [C], 6 mm [T3-1], 8 mm [T3-2], 10 mm [T3-3]) was divided by half (9-11 birds per group) to assign the antioxidant treatment (Serum [S] or MitoTEMPO [MT]). This was administered by subcutaneous injections in the skin of the back. MT-treated birds received mitoTEMPO at 1.6 mg/ml in 50 µl saline. S-treated birds received the same saline volume only. MT-treated birds received 80 µg of mitoTEMPO every other day to a total of seven doses (2.67 mg/Kg/day). First injections were performed two days after the surgery to allow birds’ recovery and ended 14 days after. The mitoTEMPO dosage was chosen from results described in mice (Vendrov *et al*. 2015), from a precedent pilot study in zebra finches, and also from an experiment in red crossbills (see also SM).

Blood samples, digital pictures of the bill, and body mass measures were taken five days before the implant date and again on the last day of the experiment. Blood samples were stored in a cold box and centrifuged 10 min at 12,000 rpm within 8h of sampling. Plasma was stored at −80°C until analyses. Body condition (i.e. std. residuals of tarsus length on body mass) was similarly distributed among the treatments and its interaction (all χ^2^ tests: p > 0.10). The treatments were randomly distributed across the aviary (cage rows and columns). Nonetheless, the two birds in each cage belonged to the same antioxidant treatment. The identity of the cage was, anyway, included as a random term in all statistical models to control for pseudo-replication. One bird (T3-1 and mitoTEMPO-treated) died by unknown reasons the day after the start of the experiment and was removed from the dataset.

Finally, the occurrence of a body feather moult was visually established at the end of the experiment by the same observer (AC). Birds were classified as engaged or not in moult (49 vs 36%, respectively).

### Thyroid hormone analyses

T3 levels were assessed from one plasma aliquot obtained in the last blood sampling event. Hormone values were measured by means of commercial species-independent ELISA kits (Arbor Assays, Ann Arbor, MI; ref. K056-H1). The analyses were made twice per sample in three sessions (intra- and inter-assay CVs = 10.2 and 14.8 %, respectively; see also SM).

### Respiratory frequency

With the aim of validating the hypermetabolic effect of thyroid hormones (e.g. Harper & Seifert 2008), the respiratory frequency was measured just before each blood sampling event by the same observer (AC). The number of breast movements in 90 seconds was counted by handing each bird face up at the left hand, with the head between the middle and forefinger (Fucikova *et al*. 2009).

### Plasma carotenoids

Total carotenoid levels in plasma were determined at the start and end of the experiment by means of spectrophotometry. Samples were diluted in ethanol, centrifuged and supernatant absorbance measured at 450 nm. The concentrations were calculated from a lutein standard curve (full description in SM). This protocol was modified from Hargitai *et al*. (2009).

### Gene expression

At the end of the experiment, a small layer of the upper surface of the upper mandible (1mm^3^ approx.) was taken with a small scalpel. The wound was disinfected and covered with blastoestimulina cream ® (Almirall labs, Spain; composed by *Centella asiatica* extract plus neomycin). All birds fully recovered in less than five weeks. The biopsied tissue was immediately introduced in RNAlater at 1:20 volume approximately and stored frozen (−20°C) until the analyses. Total RNA was extracted using the RNeasy Mini Kit (Qiagen). Residual genomic DNA carry-over was removed using the DNase treatment from the same kit. Complementary DNA (cDNA) was prepared from total RNA (∼1 µg) using the GRS cDNA Synthesis Kit (GRiSP). Quantitative real-time PCR reactions were performed on the *CYP2J19* gene (target) on cDNA. Reactions were performed by using iTaq Universal SYBR Green Supermix (Bio-Rad Laboratories) in a CFX96 Touch Real-Time PCR Detection System. β-actin was used as housekeeping (control) gene for normalizing expression levels. Primers for both *CYP2J19* and β-actin were taken from a previous study (i.e. Mundy *et al*. 2016). Mean cycle threshold (Ct) values of both genes were obtained from triplicated measures (both Lessells & Boag 1987’ *r* values = 0.99). Expression values of target genes are traditionally corrected to the expression of the control gene using a ΔCt approach, but this assumes that the expression of the control gene is kept constant across conditions. Since our experiment is likely to influence overall homeostasis, and therefore impact control gene expression, we obtained normalized Ct values using the method of Cui *et al*. (2015): normalized value = target_Ct value – (*b* x control_Ct value) where *b* is the regression coefficient of the linear regression of mean *CYP2J19* Ct values on mean β-actin Ct values. This normalization removes biases produced when the housekeeping Ct values correlate to Δ-Ct (i.e. *CYP2J19* Ct minus β-actin Ct; i.e. Cui *et al*. 2015).

### Colour measurements

Bill redness was determined by means of digital photography (Nikon® D300; full description in SM). Briefly, each bird was placed laterally and a picture of one side of the head was taken. Digital photographs were standardized and analysed using the recently developed ‘SpotEgg’ software (Gómez & Liñán-Cembrano 2017), an image-processing tool for automatized analysis of avian colouration that solves the need for linearizing the camera’s response to subtle changes in light intensity (Stevens *et al*. 2007). The mean red, green and blue (RGB) values measured from the lateral area of the bill (upper and lower mandibles) were used to obtain a hue value from the Foley & van Dam (1982) algorithm (see also Cantarero & Alonso-Alvarez 2017). Repeatability (Lessells & Boag 1987) calculated on a set of digital photographs measured twice (*n* = 30) was *r* = 0.99, *p* < 0.001. Since a low hue means a redder colour, the hue value was reversed (multiplied by −1 and adding 11 to attain positive values) to obtain a “redness” variable (also SM).

### Statistical analyses

Generalized linear mixed models (GLMMs) were used to test the treatment effects on the final values of T3, respiratory frequency, body mass difference, gene expression levels and redness as dependent variables in separated models (PROC MIXED in SAS version 9.4 software). The body mass difference (g) was calculated by subtracting the final or intermediate measure (i.e. five days before the end of the experiment) to the initial body mass (i.e. at the start of the experiment). The moult occurrence was tested by a binomial model with logit link function (PROC GLIMMIX in SAS). The hormonal treatment (a four-level factor), the antioxidant treatment (two levels), and the location of the bird into the cage (right vs. left side), as well as their interactions, were all tested in every model (also SM). The cage identity was included as a random factor and was non-significant (all *p*-values > 0.20), except in the bill redness models (*p* < 0.01). Similarly, the laboratory session in T3 and carotenoid analyses (microplate identity) were included as another random factor in models testing T3 and carotenoid plasma values, respectively, and they were also non-significant (three and four-level factor, respectively; all *p-values* > 0.22). In any event, random factors were maintained in all the models for coherence (see e.g. Bolker *et al*. 2009).

Some covariates were tested. In those models testing the body mass difference, the tarsus length was added to test size-independent variability. Bill brightness was included in the bill redness model to avoid the influence of the total amount (intensity) of light reflected from the measured surface (McGraw 2006). This allows discarding the influence of tissue structure (i.e. not pigmentation) on redness values (McGraw 2006). The bill area used to measure the redness was also tested.

The initial value of the dependent variable was additionally included as a covariate to avoid subtle initial biases. In this regard, the initial redness was calculated as the standardized residuals obtained from a model including the random factor cage identity as well as bill brightness and surface as covariates (all significant terms, *p* < 0.02). All the models were explored by backward and forward stepwise procedures, removing or including terms (*p* > 0.05 or *p* ≤ 0.05, respectively) to obtain the best fitted one. The lowest AIC criterion also agreed with this procedure. LSD *post hoc*s were used for pairwise comparisons. Plasma T3 and carotenoid values were log-transformed to attain normality. Satterthwaite DFs were used. Least square means ± standard error (LSM ± SE hereon) from mixed models are reported, excepting for moult occurrence were raw proportions are given.

## Results

### Thyroid hormone levels

The mitoTEMPO treatment or its interaction with T3 treatment did not report significant effects on plasma T3 values (both removed at *p* > 0.78). The backward procedure led to a non-significant T3 treatment factor (i.e. *F*_3,63.3_=1.50, *p* = 0.184). Nonetheless, T3-2 and T3-3 showed a trend to significantly higher values of plasma T3 than controls (LSD pairwise comparisons: *p* = 0.073 and 0.059, respectively; Figure 1A). When comparing controls versus the other birds grouped in a single group, a significant effect was detected (*F*_1,67.3_=4.07, *p* =0.048; LSM ± SE: 6.66 ± 1.01 and 9.04 ± 0.60 ng/mL, respectively).

**Figure 1.**
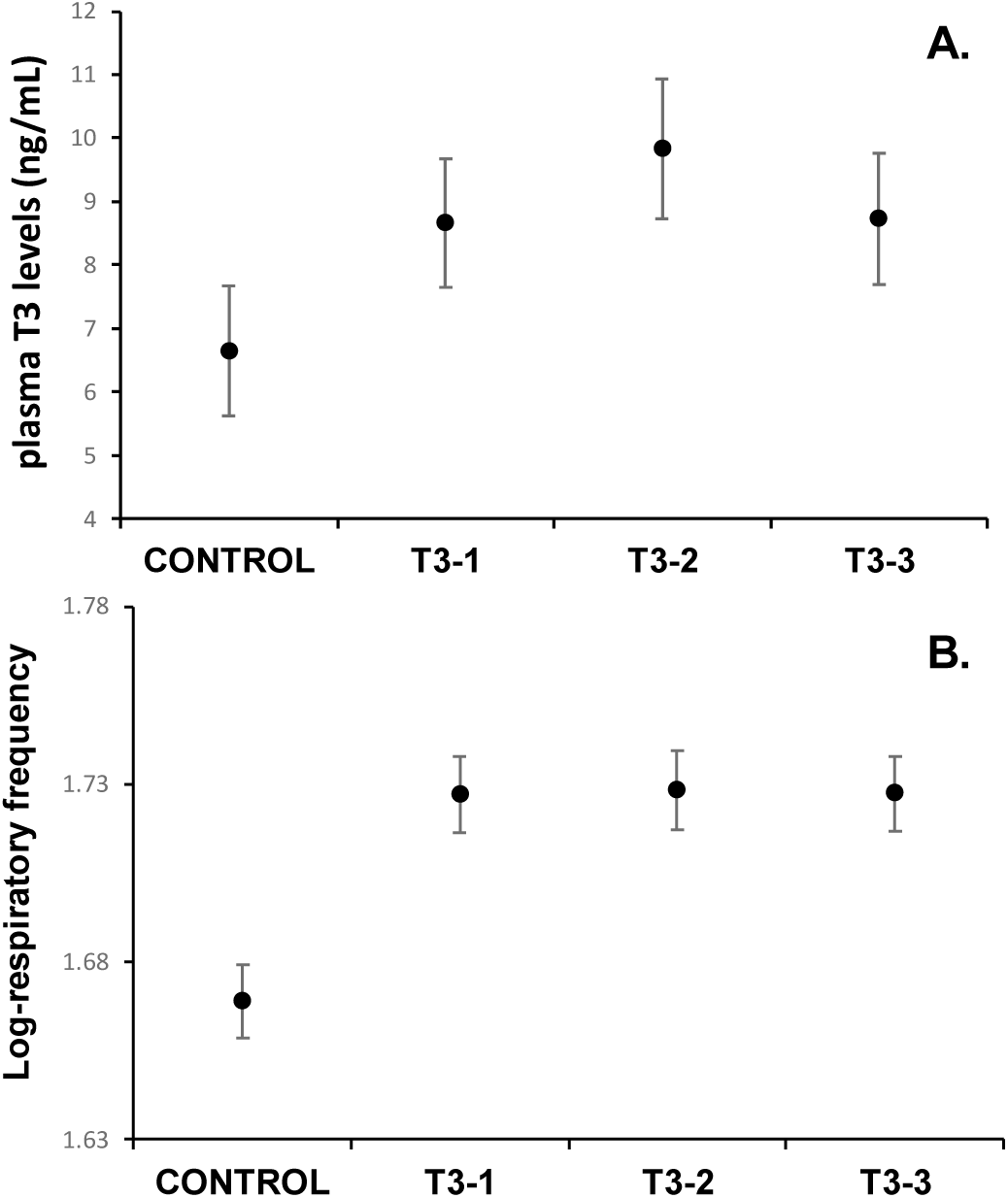
Plasma T3 levels (A) and respiratory frequency (B) at the end of the experiment depending on the length of the implant filled with T3 (T3-1: 6 mm, T3-2: 8 mm and T3-3: 12 mm). LSM ± SEs from the mixed model.

### Respiratory frequency

In the model testing respiratory frequency, only the T3 treatment effect remained (*F_3,70.7_* = 8.64, *p* < 0.001; initial value: *F_1,74.1_*= 19.09, *p* < 0.001). The LSD tests comparing the control group with each T3 treatment always reported *p* < 0.001 (Figure 1B; other pairwise comparisons *p* > 0.62).

### Moult occurrence

The T3 treatment affected the moult (*F*_3,81_= 7.28, *p* <0.001) — birds with higher doses being more frequently engaged in moult (5, 55, 85 and 91% respectively, from controls to higher dosages). Pairwise comparisons were always *p* < 0.048, excepting at the two highest levels (*p* = 0.598). No other factor remained in the model (all *p* > 0.25).

### Body mass difference

At the intermediate measure, the interaction between both treatments was non-significant (*p* = 0.301). In the best-fitted model, the antioxidant treatment reported a significant effect (Table 1). MitoTEMPO-treated birds lost less mass than controls (LSM± SE: −0.96 ± 0.09 and −1.23 ± 0.09 g, respectively). T3 treatment was also a significant factor (Table 1) because birds in any hormone-treated group lost more mass than controls (all comparisons: *p* < 0.001; Figure 2A). The T3-1 vs T3-3 comparison was also significant (*p* = 0.027). When the final body mass was tested, the treatment interaction reported (*p* = 0.085). Only mitoTEMPO-treated birds not exposed to exogenous T3 increased body mass (Figure 2B), with their weight differing from all other groups (all *p*-values < 0.028). Serum-only treated birds also differed or tended to significantly differ from birds in T3-2 and T3-3 dosages (*p*-range: 0.018-0.066; other *p* > 0.10). If the interaction is removed, only the hormone treatment remained (*F_3,68.2_* = 10.58, *p* < 0.001), controls differing from any other group (all *p-values* < 0.001; other *p-*values > 0.15; see Table S1 and Figure S1).

**Figure 2.**
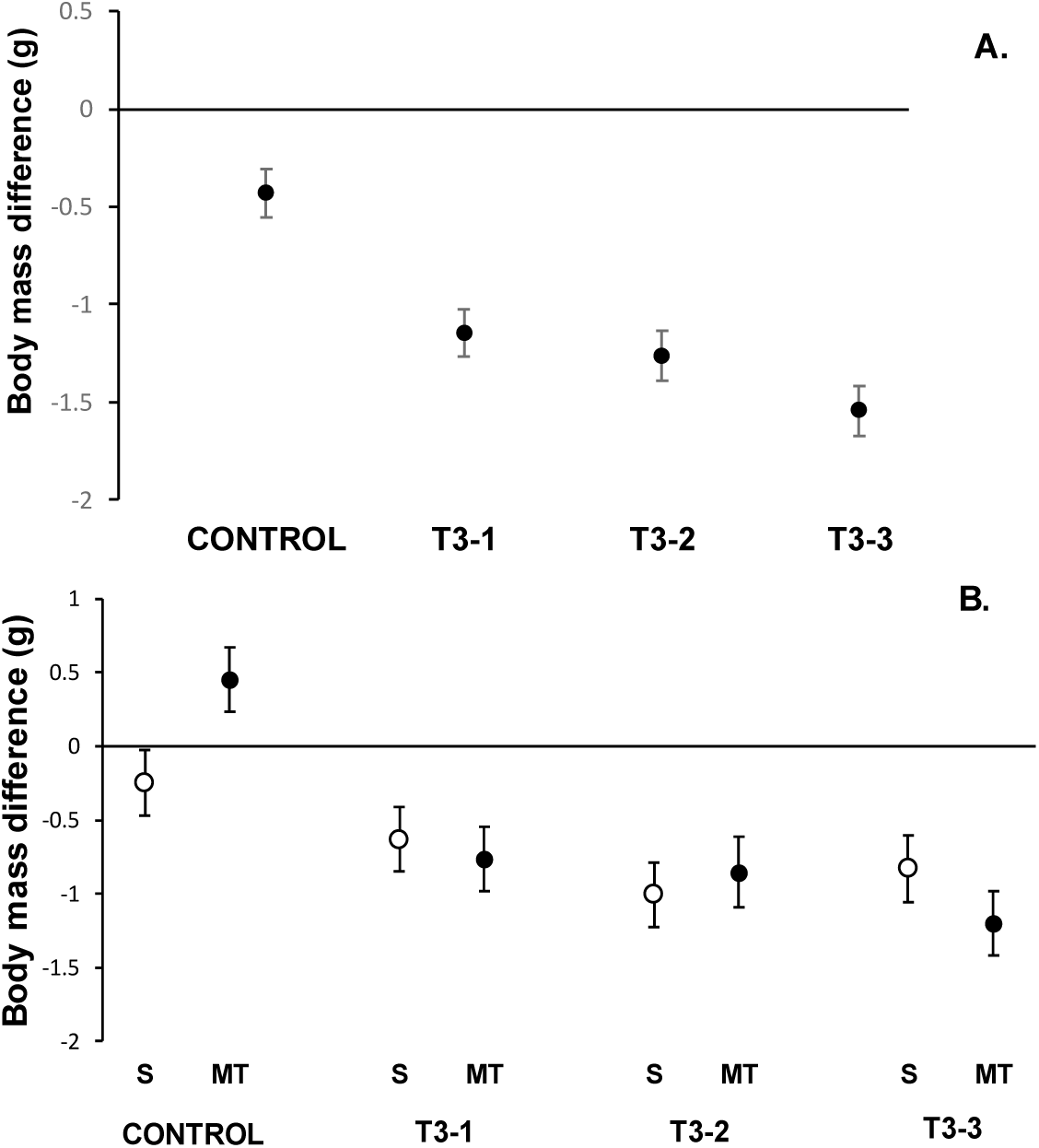
Difference between intermediate (A) and final (B) body masses regard to initial pre-experimental values depending on T3 dosage (T3-1: 6mm, T3-2: 8mm and T3-3: 12mm) and antioxidant treatment (S: serum; MT: mitoTEMPO). LSM ± SEs from mixed models controlling for body size (tarsus length) variability.

**Table 1.** Best fitted models testing body mass variability at two different sampling times (intermediate and final, see main text).

| *Intermediate body mass* | **Slope ± SE** | **F** | **df** | **p** |
| --- | --- | --- | --- | --- |
| MitoTEMPO treatment | - | 4.38 | 1,41 | 0.043 |
| T3 treatment | - | 15.09 | 3,65.7 | < 0.001 |
| Initial body mass | -0.175 ± 0.043 | 16.52 | 1,76.9 | < 0.001 |
| Tarsus length | 0.244 ± 0.116 | 4.45 | 1,74.6 | 0.038 |
| *Final body mass* | **Slope ± SE** | **F** | **df** | **p** |
| MitoTEMPO treatment | - | 39.5 | 0.29 | 0.595 |
| T3 treatment | - | 65.3 | 2.30 | 0.085 |
| Initial body mass | -0.173 ± 0.054 | 73.9 | 10.13 | 0.002 |
| Tarsus length | 0.430 ± 0.149 | 71.3 | 8.35 | 0.005 |

### Circulating carotenoids

No term reported a significant effect in the model testing plasma carotenoid levels (all *p*-values > 0.22).

### *CYP2J19* expression

The treatments reported a clear highly significant interaction (Table 2). The resulting figure (Figure 3A) resembled that of bill redness at least in the two highest T3 dosages (Figure 3B). MT vs S comparisons at T3-0 (controls), T3-2 and T3-3 hormonal dosages reported *p* = 0.035, 0.032 and 0.055, respectively. A global view suggests a U-shape relationship with hormone dosages among S-injected birds, whereas an inverted U appears among mitoTEMPO-injected animals. Thus, S-injected birds reported lower *CYP2J19* expression at the medium-sized (T3-2) dosage compared to hormone controls (*p* = 0.014; other comparisons among S-injected birds: *p* > 0.082). In mitoTEMPO-treated birds, the control vs T3-2 reported *p* = 0.066, suggesting increased gene expression (Figure 3B). The value then declined at the highest T3 dosage (T3-1 vs T3-3 and T3-2 vs T3-3: both *p-values* < 0.027; other comparisons: *p*-values > 0.17). It is worth noting that when circulating carotenoids at bill sampling time were added as a covariate, they were positively correlated to gene expression (slope ± SE: 0.059 ± 0.029; *F*_1,72.5_ = 4.18, *p* = 0.045), but this did not alter the treatment interaction *(p* = 0.013).

**Figure 3.**
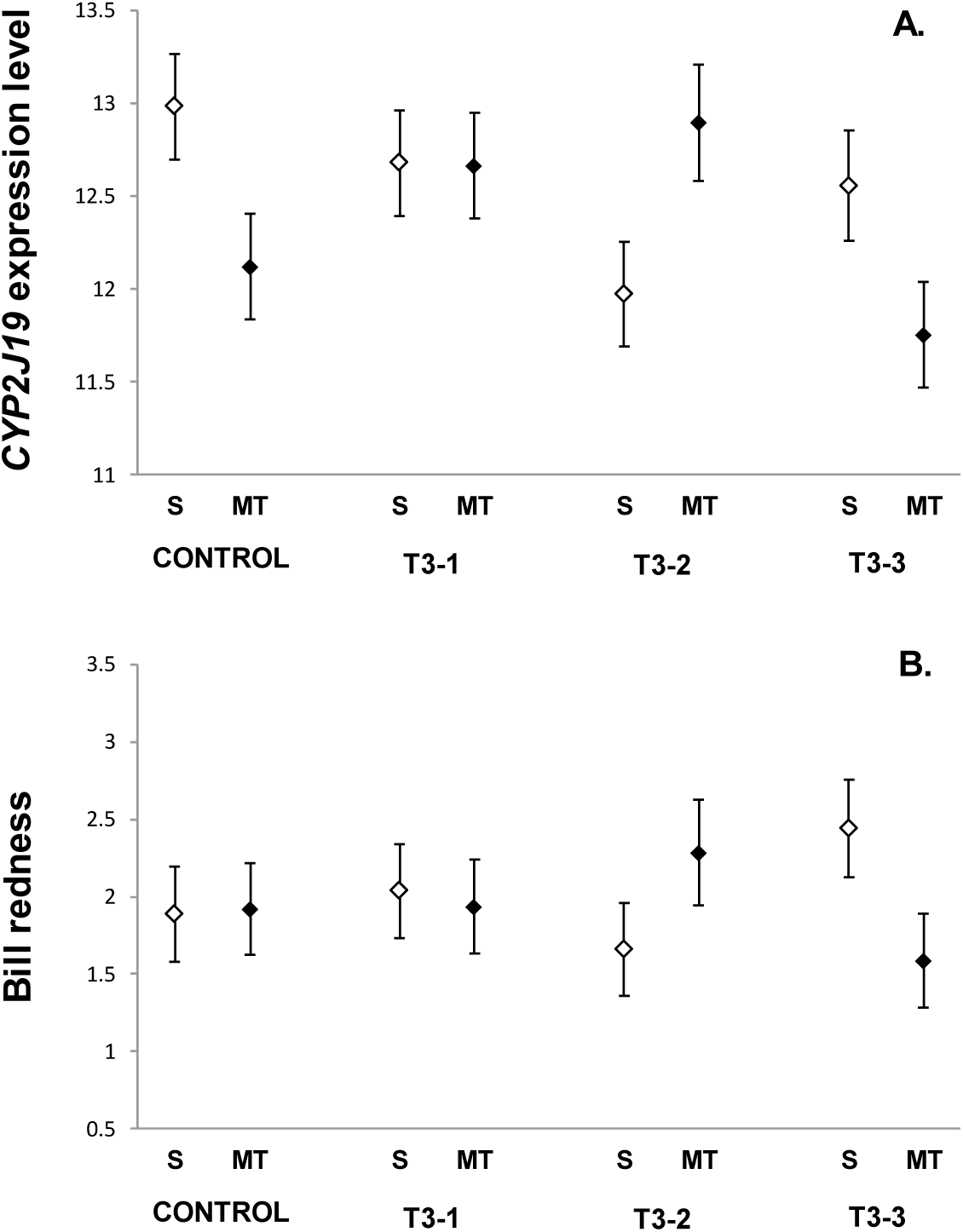
Treatment effects on the *CYP2J19* expression level (A) and bill redness (B). See the description of variables in Methods. (S: serum only; MT: MitoTEMPO-injected birds: T3-1: 6mm, T3-2: 8mm and T3-3: 12mm). LSM ± SE from mixed models.

**Table 2.** Best fitted models testing the impact of the mitoTEMPO and T3 treatments on male zebra finch bill *CYP2J19* expression and redness.

| *CYP2J19* | Slope ± SE | *F* | df | *p* |
| --- | --- | --- | --- | --- |
| MitoTEMPO treatment | - | 0.71 | 1,42.2 | 0.405 |
| T3 treatment | - | 1.20 | 3,65.6 | 0.318 |
| MitoTEMPO x T3 treatments | - | 4.15 | 3,67.5 | 0.009 |
| *Bill redness* | Slope ± SE | *F* | df | *p* |
| MitoTEMPO treatment | - | 0.08 | 1,38.7 | 0.777 |
| T3 treatment | - | 0.08 | 3,53.3 | 0.973 |
| MitoTEMPO x T3 treatments | - | 2.42 | 3,58.6 | 0.075 |
| Location into the cage | - | 9.49 | 1,29.4 | 0.004 |
| Measured area (mm<sup>2</sup>) | 0.055 ± 0.027 | 4.22 | 1,50.6 | 0.045 |
| Initial bill redness (residuals) | 0.355 ± 0.161 | 4.85 | 1,37.5 | 0.034 |
| Bill brightness | -6.919 ± 1.007 | 47.18 | 1,55.8 | <.0001 |

### Bill redness

The interaction between both treatments reported a trend to significance (*p* = 0.075; Table 2; also SM for descriptions on other significant terms). This was driven by differences between antioxidant treatments in the two groups with the larger implants (Figure 3A). In serum-injected birds, bill redness increased between T3-2 and T3-3 doses (*p* =0.024), whereas the opposite pattern was suggested among mitoTEMPO-treated birds (but *p* = 0.135). Moreover, in T3-3-treated individuals, the mitoTEMPO effect showed a trend to significance (*p* = 0.062), here reversing the positive effect of the hormone on redness.

Finally, when *CYP2J19* expression level is included as another covariate in the redness model, the interaction became clearly non-significant (*p* = 0.209), both treatments being subsequently removed (both *p* > 0.80). This covariate then showed a significant positive relationship with redness (i.e. *F*_1,78_ = 5.52, *p* = 0.021; slope ± SE: 0.242 ± 0.103). The regression on raw data was also significant (*r* = 0.26, *p* = 0.017, see Figure 4 and also S3).

**Figure 4.**
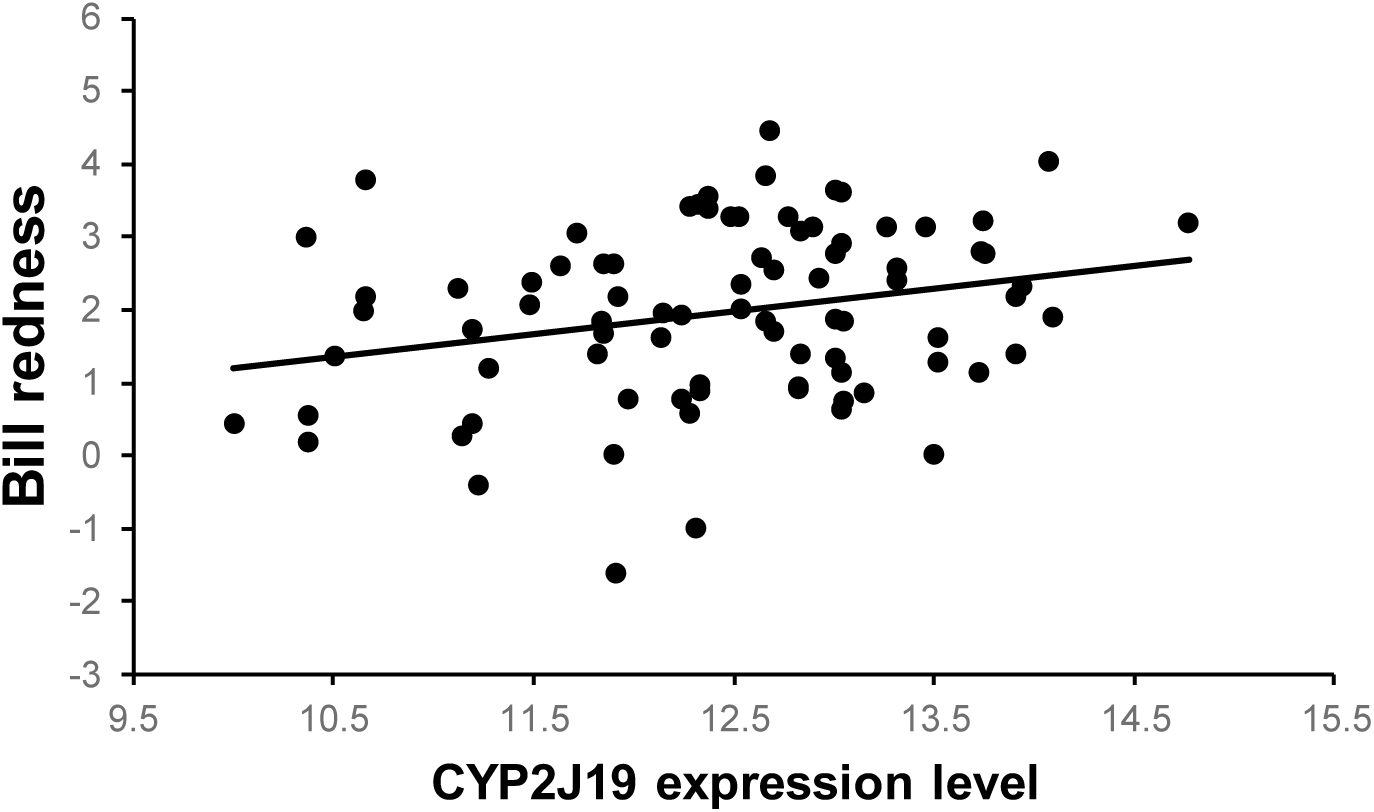
Relationship between male zebra finch bill redness and the expression level of the candidate enzyme for ketocarotenoid synthesis in that ornament (slope ± SD: 0.310 ± 0.127). Raw data with redness as the reversed hue value (see Methods).

## Discussion

We have shown that *CYP2J19* expression levels at the bill epidermis positively correlates to the expression of a red carotenoid-based trait and that this link is affected by a mitochondrial antioxidant and a thyroid hormone. T3-treated birds as a single group showed higher hormone levels than controls, as well as higher respiratory frequency and body mass loss, being also engaged in moult. These effects are well-supported by avian literature (Elliott *et al*. 2013; Welcker *et al*. 2013; McNabb & Darras 2015; Perez *et al*. 2018; also SM) and indicate that implants were effective. Moreover, birds with the longest and shortest T3 implants differed in body mass loss, which suggests that the T3 dosage was also relevant. Bill redness and *CYP2J19* expression variability support this view. We should, hence, assume that the lack of significant differences in circulating hormone levels among T3-1 to T3-3 groups was probably due to feedback regulation (Leung, Taylor & Van Iderstine 1985; Hull *et al*. 1995; Perez *et al*. 2018), with the effects on target tissues, anyway, differing.

First, we should address the question of why mitoTEMPO-treated birds lost less body mass than other individuals. It contradicts studies where rodents fed with a high caloric diet did not gain mass when treated with this compound (Jeong *et al*. 2016; Gutiérrez-Tenorio *et al*. 2017). Here, mitoTEMPO seems to buffer the body mass loss probably derived from handling stress (see McGraw, Lee & Lewin 2011). In fact, contrarily to typical fatness induced by reduced locomotor activity under captivity, most of our birds lost body mass (Figure 3). Anyway, the addition of body mass change (%) as a covariate to the redness model, even when negatively correlated (birds gaining mass losing redness), did not alter our results (Table S2 and Figure S2). Moreover, this body mass change did not correlate with *CYP2J19* expression (covariate always *p* > 0.80).

In regard to *CYP2J19*, we must first highlight that mitoTEMPO and T3 treatments did not influence circulating carotenoid levels, and the addition of plasma carotenoid values as a covariate to the CYP2J19 model, even when correlated, did not alter the results. Therefore, the experimental effects appear to be, at least partially, independent of changes in substrate availability (yellow carotenoids; McGraw & Toomey 2010). This may support recent ideas suggesting that carotenoid-based signalling is not dependent on resource allocation trade-offs (Koch & Hill 2018) as often defended (e.g. Alonso-Alvarez *et al*. 2008; García-de Blas, Mateo & Alonso-Alvarez 2016).

Moreover, we detected a significant interaction between mitoTEMPO and T3 treatments on *CYP2J19* expression, but the results did not follow our initial predictions (Figure 4). Contrarily to our expectations, mitoTEMPO downregulated *CYP2J19* expression among hormone controls, although this was not reflected in bill redness variability (see also below). Such a ketolase downregulation apparently contradicts the effects of another mito-targeted molecule (mitoQ) that improved zebra finch bill redness (Cantarero & Alonso-Alvarez 2017). We argue that this is a consequence of subtle differences in the action mechanisms of mitoQ and mitoTEMPO. The first is a synthetic ubiquinone that improves electron transfer at the IMM, whereas mitoTEMPO is a superoxide dismutase (SOD) mimetic that favours ubiquinone recycling by reducing superoxide production (Murphy & Smith 2007; Trnka *et al*. 2008; Dikalova *et al*. 2010). Another difference is the shorther length of the alkyl chain of mitoTEMPO compared to mitoQ. This is likely to have improved the antioxidant action, avoiding disruption of the mitochondrial membrane and higher superoxide production (Reily *et al*. 2013; Trnka, Elkalaf & Anděl 2015; Gottwald *et al*. 2018). In this regard, a comparison of the effects of mitoQ and a very similar compound (mitoTEMPOL) in human cancer cell lines revealed lower levels of superoxide generation in those cells treated with the second compound (Pokrzywinski *et al*. 2016). We thus hypothesize that mitoTEMPO induced a strong reduction in superoxide levels (see Dikalova *et al*. 2010) that could have disrupted cell redox signalling mechanisms linked to ketolase gene expression. We note that superoxide is a well-known redox signal affecting the expression of many genes (reviewed in Hurd & Murphy 2009; Weidinger & Kozlov 2015). In support of this, it has been shown that mitoTEMPO can induce downregulation of human inflammatory and cancer-related genes by interfering redox signalling pathways as a result of a decline in mitochondrial superoxide generation (Nazarewicz *et al*. 2013; McCarthy & Kenny 2016).

The inhibitory effect of mitoTEMPO on *CYP2J19* expression also contradicts a recent study where male captive red crossbills treated with the same compound and dosage have regrown redder feathers after plucking (Cantarero *et al*. 2019 preprint). Interestingly, in that study, the effect was only detected among the reddest birds at the start of the experiment (supposedly the high-quality animals; Cantarero *et al*. 2019 preprint). This strongly points to other factors controlling mitoTEMPO action on ketolase activity. Perhaps the answer comes from the mitoTEMPO x T3 interaction here showed.

Thus, the mitoTEMPO-induced *CYP2J19* downregulation showed in hormone controls disappeared from T3-1 to T3-2 (Figure 4A). The hormone treatment effect on antioxidant controls, in fact, inhibited *CYP2J19* expression throughout the same dosage range. As previously mentioned, in mammals, but also birds, thyroid hormones have been linked to high ROS generation and oxidative stress (reviewed in Venditti & Meo 2006; Rey *et al*. 2013; Venditti *et al*. 2015) due to increased oxidative metabolism (i.e. oxygen consumption rate) (e. g. Hulbert 2000), and we know that ROS can inhibit CYP levels and activity (see El-Kadi *et al*. 2000 for an example of oxidative stress on P450 activity). The presence of the mito-targeted antioxidant mitoTEMPO at T3-2 dosage reversed the inhibitory effect of T3 and even induced higher ketolase expression compared to hormone controls treated with the antioxidant. Subsequently, at the highest T3 dosage, the birds seem to be able to trigger some compensatory mechanism as *CYP2J19* expression apparently increased (Figure 4A).

Literature from mammalian models supports the capacity of thyroid hormones to mount compensatory/protective responses against its own pro-oxidant effects (e.g. reviewed in Villanueva, Alva-Sánchez & Pacheco-Rosado 2013; Goharbari, Shadboorestan & Abdollahi 2016). For example, in rats, hyperthyroidism leads to increased activity of thioredoxin/peroxiredoxin enzymes transforming hydrogen peroxide (derived from superoxide) to water (Venditti *et al*. 2015). T3 can also upregulate genes involved in mitochondrial biogenesis, thus reducing mitochondrial damage accumulation (Weitzel & Alexander Iwen 2011) or directly upregulating gene expression of different p450 enzymes (Brtko & Dvorak 2011; Tee *et al*. 2011). In birds, T3-treated chicken increased SOD activity compared to controls, which apparently avoided oxidative damage (Lin, Decuypere & Buyse 2008). In the same line, T3-treated Muscovy ducklings (*Cairina moschata*) upregulated the gene expression of uncoupling proteins (UCPs) located at the IMM, which are involved in decoupling cell respiration from ATP synthesis allowing a decrease in superoxide generation (e.g. Rey *et al*. 2010). In any event, mitoTEMPO seems to be able to disable that compensatory mechanism (see Figure 4 at T3-3), perhaps by interfering in redox signalling such as suggested for the hormone control group (above).

Although *CYP2J19* expression levels and redness showed the same pattern of change at the two highest T3 dosages (Figure 4), the correlation between both variables is far from perfect (Figure 5). Thus, the mitoTEMPO-induced *CYP2J19* downregulation did not lead to paler bills among hormone controls. This perhaps reveals a delay in the mitoTEMPO effect as colouration could not only depend on the number of ketolase copies but on ketolase activity. Moreover, colouration effects could also have been mediated by post-transcriptional regulation attenuating the impairing effect on colouration (e.g. Smutny, Mani & Pavek 2013). The fact that *CYP2J19* expression and redness variabilities varied in concert at the highest T3 dosages may suggest that the mitoTEMPO impact on the phenotype was quicker under the hypermetabolic effects of the thyroid hormone.

Overall, our results reveal an interacting effect of mitochondrial antioxidant metabolism and thyroid hormones that suggests that certain levels of superoxide are needed to maintain efficient carotenoid biotransformation (see also García-de Blas, Mateo & Alonso-Alvarez 2016). Understanding this mechanism would require future correlational and experimental approaches. This includes determining mitochondrial superoxide production changes as well as variability in ketocarotenoid concentrations at the ornament tissue. Nonetheless, this is the first study showing a positive significant correlation between *CYP2J19* expression level and red colouration in any species. Moreover, mitoTEMPO effects give additional support to the hypothesis that the mitochondrion is involved in colour-based sexual signalling (Johnson & Hill 2013; Cantarero & Alonso-Alvarez 2017), also supporting that red ketocarotenoid-based ornaments could act as indices revealing the individual capacity to efficiently perform cell respiration under a sexual selection scenario (Hill 2011; Hill & Johnson 2012). Finally, thyroid effects on avian carotenoid-based colouration have virtually been ignored. We have only been able to find a single study (Schereschewsky 1929), where male bullfinches (*Pyrrhula pyrrhula*) supplied with a thyroid extract moulted paler plumages, which may support some of our findings. Our results thus open a landscape for future studies.

## Acknowledgements

We thank Luisa Amo for providing us with some of the birds in the experiment and Lucía Arregui and Diego Gil for assistance in thyroid hormone quantification. Financial support was obtained from the project CGL2015-69338-C2-2-P (MINECO). AC is currently supported by a postdoctoral fellowship from Fundación Ramón Areces. The experiment was approved by the Bioethics Committee of CSIC and Junta de Castilla La Mancha (JCCM) government (ref number 16-2017).

## Supplementary Material

### Additional methodological details

The birds were housed at the Fundación para la Investigación en Etología y Biodiversidad (http://es.fiebfoundation.org). All birds were given *ad libitum* access to food (commercial pelleted food; KIKI®, Spain), water and grit. The temperature (mean ± range 22° ± 1°C) and light daily cycle (16L : 8D) were controlled.

A small incision with a scalpel was made in the skin of birds to introduce the implants, and surgical glue (Cicastick®) of veterinary use was used to close the wounds, though suture (two 17-70 cm stitches of Braun®, Model Novosyn Violet, 6/0 HR) was additionally used.

The mitoTEMPO dosage was chosen from a pilot study involving 10 male zebra finches randomly assigned to different concentrations (0, 0.334, 0.668, 1.335 and 2.67 mg/Kg/day) subcutaneously injected in 50 µl saline every other day for two weeks. The highest level (2.67 mg/Kg/day; 3mM) was chosen as we did not find a significant correlation between dose and body mass change (%; Spearman’s *r* = 0.320, *p* = 0.367), suggesting no health impairment. Moreover, no evident toxicity symptoms (behaviour changes, fatigue, lack of alertness) were detected and the change (%) in bill redness increased with dosage (Spearman’s *r* = 0.82, *p* = 0.01; see also Cantarero & Alonso-Alvarez 2017 for colour analysis methods)(Cantarero & Alonso-Alvarez 2017). The final dosage was, nonetheless, higher than that reported in mice (1.5 mg/Kg/day; Vendrov *et al*. 2015) where mitoTEMPO decreased mitochondrial free radical production. In the cited study, however, the antioxidant was daily administered for 84 days (here 14 days).

The origin of the birds (three commercial suppliers and one experimental population [i.e. IREC-CSIC; Ciudad Real, Spain]) was also balanced among antioxidant treatments (*χ*^2^=2.73, df=3, *p* = 0.435), T3 implant size (*χ*^2^= 3.86, df = 9, *p* = 0.920) and among the eight combinations (antioxidant x hormone groups: *χ*^2^= 10.81, df = 21, *p* = 0.966).

### Implant rejection events

The explanation to implant rejection experienced by some birds was unclear. Although we initially suspected that it could have been due to the implant sealing method, trials with different alternatives (glues or suturing the silicone tube ends) had similar rejection rates. The same Silastic® (Dow Corning) implant type has been used in avian endocrinology studies for decades (Wingfield 1984 and many articles from that researcher; see recently e.g. Noguera, Kim & Velando 2017). We suspect that silicone composition has recently subtly changed as the patented product is currently manufactured by another company (Freudenberg Medical), and we must consider that it was designed for humans/rodent models. In any event, the lack of effect of the “days with implants” covariate and also the randomized distribution of treatments among those birds losing implants suggests that the impact of implant rejection on main conclusions is likely to have been minor.

### Hormone analyses

Plasma aliquots were allowed to defrost and equilibrate for an hour at room temperature. A volume of 30 µl of plasma was transferred to labelled glass tubes. Hormone extraction was made by adding 3 ml of diethyl ether to the tubes, vortexing for 5 min and then centrifuging (5 min at 1500 rpm) in at 4°C. These tubes were maintained in a freezing bath of ethanol plus dry ice for 2 min. The etheric phase was transferred to a new clean tube, which was left to dry out in a warm bath under a fume hood (30 min at 40°C). This extraction protocol was performed twice. Extractions were re-suspended with 130 µl of steroid buffer (Arbor Assays, Ann Arbor, MI; ref. K056-H1) and vigorously vortexed for 5 min.

The measurements were performed by means of commercial ELISA kits (Arbor Assays, Ann Arbor, MI; ref. K056-H1). A microplate reader was used (Multi-detection Synergy HT; Biotek®). The standard curves provided a very good fit to standards (r^2^ > 0.99). The detection limit of the assay (80% maximum binding) was found at 0.078 ng/ml. The assays were made twice per sample (intra- and inter-assay CVs = 10.2 and 14.8 % respectively). The recoveries (%) described in another avian study using a similar ELISA kit ranged 93-113% (Elarabany, Abdallah & Said 2012). The analyses were made following the advice of Arbor Assays technicians.

Plasma T3 values in our control and T3-treated birds (mean ± SD, range: 6.64 ± 2.68, 3.83-15.05 and 9.04 ± 5.27, 2.48-29.02, respectively) where within the range reported for this species in other studies (Eng, Williams & Elliott 2013; Yamaguchi *et al*. 2017).

### Respiratory frequency

The number of breast movements in 90 seconds was counted by handing the birds face up at the left hand, with the head between the middle and forefinger. Breast movements were registered for 90 s divided into two 45 s bouts. In between these two bouts, the birds were introduced in a fabric bag while its cage mate was captured and its first bout of respiratory frequency measurement was taken (i.e. 2 min approx.). The two 45 s measurements taken to calculate the respiratory frequency value at both the beginning and end of the experiment were highly repeatable (Lessells & Boag 1987)(Lessells & Boag 1987; initial value: *r* = 0.93, final value: *r* = 0.97, both *p* <0.001). After the measurement, the birds were weighed and bleed.

### Plasma carotenoid quantification

Plasma aliquots were allowed to defrost for 20 min at room temperature. 10 µl were transferred to labelled plastic tubes containing 90μl of absolute ethanol, being then vortexed (3 min) and centrifuged (5 min at 11000g) in a cooled centrifuge (4°C) to precipitate flocculent proteins. The absorbance of the supernatant was measured at the lutein peak (450 nm) by spectrophotometric analyses, using a seven-level calibration curve obtained from serial dilution derived from a tube containing 22.5 μg of lutein in 1500 μL absolute ethanol. The protocol was modified from Hargitai et al. (2009).

### Digital photography

For each photo, the same standard grey reference and scale (ColorChecker Classic target; X-Rite, Michigan) was placed next to the bird’s head. The focus and diaphragm of the camera were manually fixed to avoid the interference of automatic functions. We have previously shown that these picture-based measurements are highly correlated with the redness measurement (i.e., red hue) obtained from portable spectrophotometers (Mougeot *et al*. 2007; Alonso-Alvarez & Galván 2011). SpotEgg software, however, allows the user to manually draw any region and provide information about its colouration, shape or other features. The measure of a large delimited area, as opposed to portable spectrophotometers that analyse colouration of reduced spots (usually 1-2 mm), makes this tool useful for evolutionary biologists aiming to capture most of the variability among individuals (Gómez & Liñán-Cembrano 2017). Accordingly, for each animal, the average of red, green, and blue (RGB) components of the lateral bill surface (upper and lower mandibles) were calculated. We used the lateral side of the bill due to the low variance of colouration at the top. We then determined hue values by means of the Foley & van Dam algorithm (1982) (see also main text).

### Additional analyses

In the model testing the difference between final and initial size-corrected body mass, the backward procedure at *p* < 0.05 reported that only the hormone treatment factor was retained (Table S1 and Figure S1).

**Table S1.** Mixed model testing the body mass difference between final and initial measures controlled for body size (tarsus length) and also initial body mass.

| *Final body mass difference* | slope± SE | *F* | df | *P* |
| --- | --- | --- | --- | --- |
| T3 treatment | - | 10.58 | 3,68.2 | < 0.001 |
| Initial body mass | -0.155 ± 0.055 | 8.13 | 1.78.8 | 0.006 |
| Tarsus length | 0.352 ± 0.149 | 5.66 | 1.77 | 0.020 |

**Figure S1.**
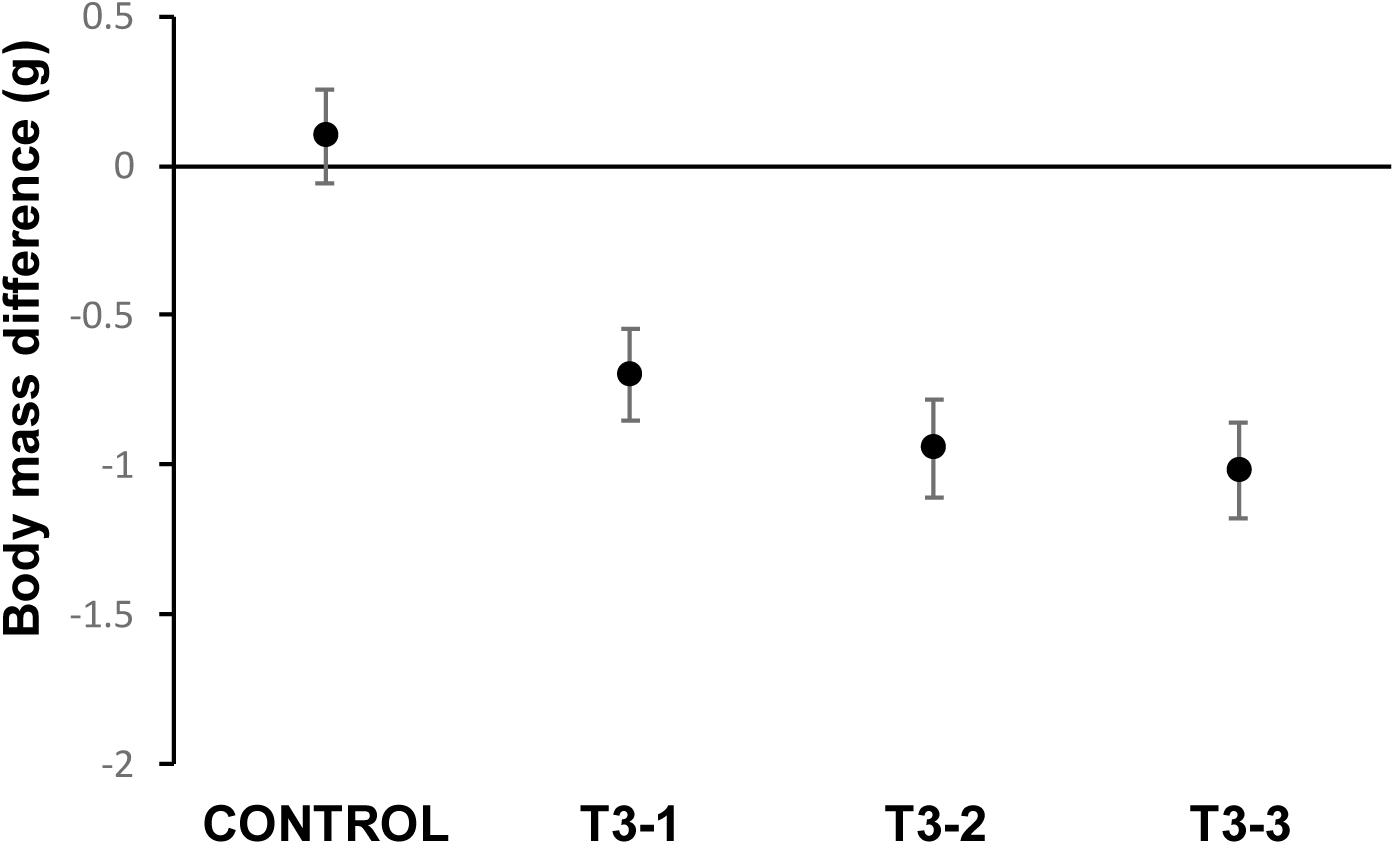
Body mass difference between final and initial measurement depending on T3 dosage (T3-1: 6mm, T3-2: 8mm and T3-3: 12mm). LSM ± SEs from mixed models.

In the model testing final bill redness, when we added body mass change during the experiment (% from initial to final measure) in order to assess hormonal effects independently of T3-mediated body mass effects, the interaction becomes significant (Table S1 and Figure S1). Here, the antioxidant effect at the highest T3 dosage is clearer (*p* = 0.032; LSM ± S.E.: mitoTEMPO: 0.424 ± 0.304; serum: 1.386 ± 0.310). The difference between the two highest T3 dose groups was again evident among antioxidant controls (*p* =0.01; LSM ± S.E.: 0.550 ± 0.297 and 1.386 ± 0.310, for T3-2 and T3-3, respectively). Among those treated with mitoTEMPO, the T3-2 vs T3-3 comparison reported a *p*-value = 0.079 (LSM ± S.E.: 1.228 ± 0.332 and 0.424 ± 0.304, for T3-2 and T3-3, respectively), thus suggesting that birds became paler at the highest dose.

**Table S2.** Model testing the impact of mitoTEMPO and T3 treatments on male bill redness when controlling for body mass change (%; bold) and other factor and covariates.

| | Slope ± S.E. | *F* | df | *p* |
| --- | --- | --- | --- | --- |
| mitoTEMPO treatment | - | 0.05 | 1, 40.2 | 0.828 |
| T3 treatment | - | 0.22 | 3, 55.5 | 0.885 |
| MitoTEMPO x T3 treatments | - | 3.40 | 3, 56.8 | 0.024 |
| Cage side | - | 7.56 | 1, 30.2 | 0.010 |
| Measurement area | 0.058 ± 0.026 | 5.18 | 1, 47.7 | 0.027 |
| Initial Redness (residuals) | 0.276 ± 0.153 | 3.25 | 1, 36.2 | 0.0797 |
| Final bill brightness | -6.999 ± 0.958 | 53.39 | 1, 52.4 | <.0001 |
| **Body mass change (%)** | **-0.042 ± 0.017** | **6.08** | **1, 48.6** | **0.017** |

**Figure S2.**
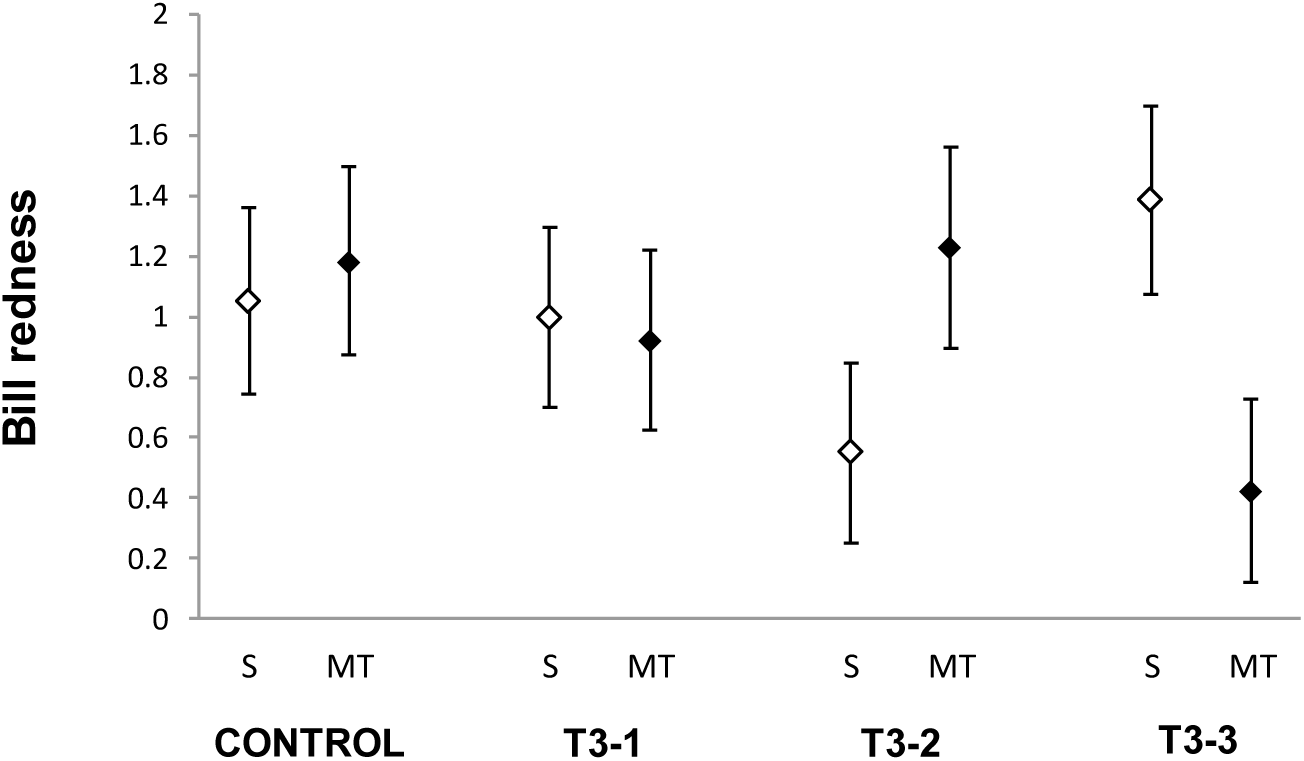
Treatment effects on the bill redness. Bill redness was the result of reversing the hue value (x −1) making data positive by adding a fixed value of 11 (see main text; S: serum only; MT: MitoTEMPO-injected birds: T3-1: 6mm, T3-2: 8mm and T3-3: 12mm). LSM ± SE from mixed models controlling for body mass change (%) during the experiment (see Results).

**Figure S3.**
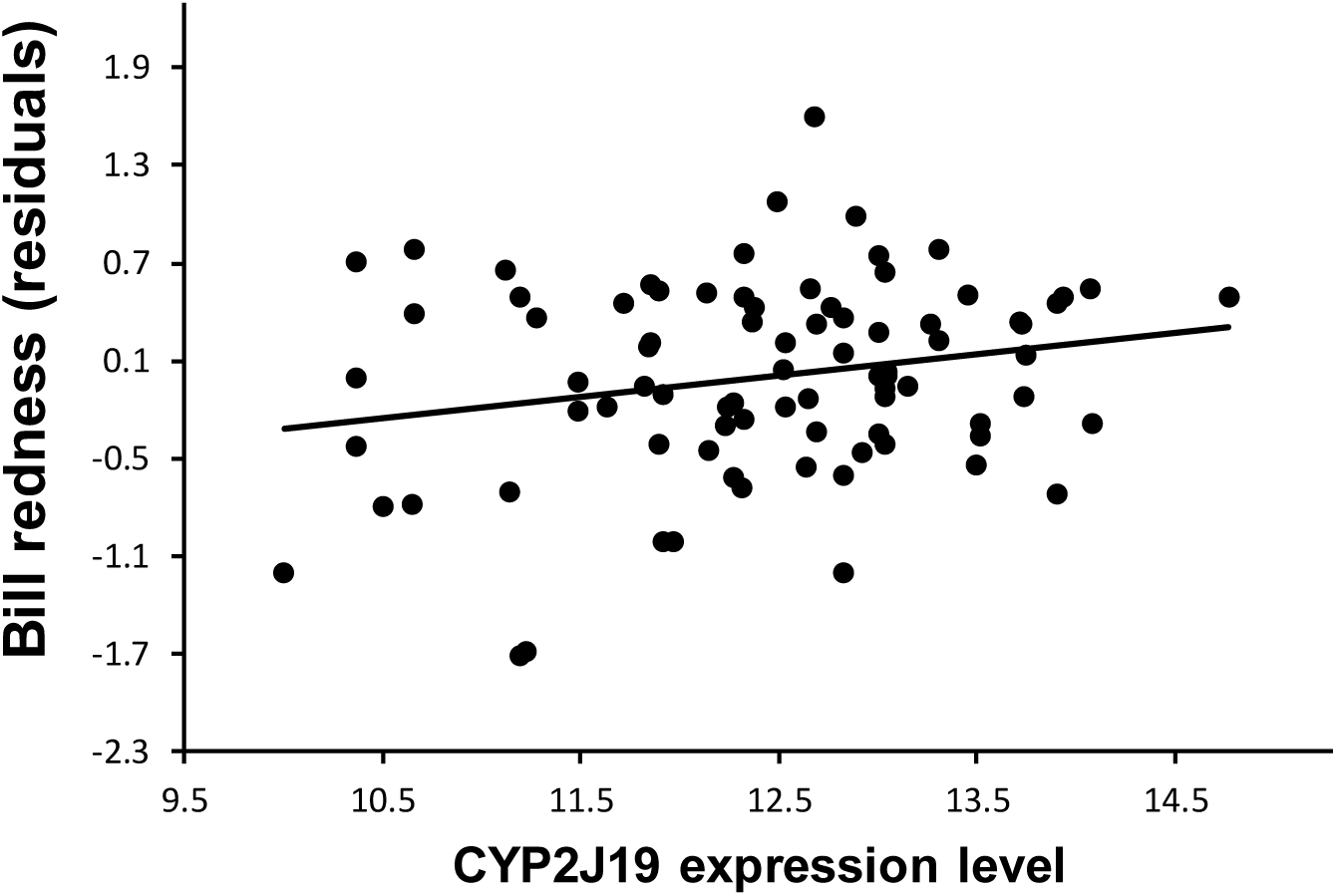
The relationship between bill redness and CYP2J19 expression level. The redness level is here controlled for any other term in the redness model (bill area, brightness, side of the cage and cage identity random factor). See Results section in the main text for the slope ± SE and other details.

### Other significant terms in the redness model

In the model testing bill redness (Table 2 in the main text), the location into the cage affected the dependent variable. The bill of those birds placed at the left side was redder than the bill of birds at the right side (LSM ± S.E.: 1.239 ± 0.157 and 0.699 ± 0.157, respectively). This could perhaps be interpreted as birds in the right side being subtly exposed to higher stress levels inhibiting red colouration production. In this regard, we must note that maintenance work daily made (water and food supply) was always made from the left to the right side of the cage. This may have exposed to birds placed in the right side to higher disturbance as both cage sides were only separated by a grille, the bird on the right side enduring more time of disturbance. This apparently subtle influence did only induce a significant effect in the case of the redness model.

Regarding other significant terms in the same model (Table 2 in the main text), the bill area was positively related to redness, i.e. larger bills were redder. We have not found a reasonable technical explanation to this effect, though it could be related to individual quality.

The initial bill redness, as well as bill brightness at the sampling time, were positively and negatively, respectively, related to final bill redness (Table 2 in the main text). This would, respectively, mean that bill redness was individually repeatable and paler bills reflected more light.

